# Theoretical Analysis of Divalent Cation Effects on Aptamer Recognition of Neurotransmitter Targets

**DOI:** 10.1101/2023.11.15.567205

**Authors:** Ali Douaki, Annina Stuber, Julian Hengsteler, Dmitry Momotenko, David M. Rogers, Walter Rocchia, Jonathan D. Hirst, Nako Nakatsuka, Denis Garoli

## Abstract

Aptamer-based sensing of small molecules such as dopamine and serotonin in the brain, requires characterization of the specific aptamer sequences in solutions mimicking the *in vivo* environment with physiological ionic concentrations. In particular, divalent cations (Mg^2+^ and Ca^2+^) present in brain fluid, have been shown to affect the conformational dynamics of aptamers upon target recognition. Thus, for biosensors that transduce aptamer structure switching as the signal response, it is critical to interrogate the influence of divalent cations on each unique aptamer sequence. Herein, we demonstrate the potential of molecular dynamics (MD) simulations to predict the behaviour of dopamine and serotonin aptamers on sensor surfaces. The simulations enable molecular-level visualization of aptamer conformational changes that, in some cases, are significantly influenced by divalent cations. The correlations of theoretical simulations with experimental findings validate the potential for MD simulations to predict aptamer-specific behaviors on biosensors.

---

Nucleic acid-based aptamers are single-stranded, short DNA or RNA sequences, isolated *in vitro* from a library of synthetic oligonucleotides using a combinatorial technique termed systematic evolution of ligands by exponential enrichment (SELEX).^1,2^ Oligonucleotides from the library are selected based on affinity and selectivity toward a specific target analyte of interest. In recent years, significant progress has been made in the development of biosensors with aptamer-coupled transducers.^3,4^ Among the diverse biosensing platforms that can be integrated with aptamers, field-effect transistors (FETs) enable direct electronic target detection with high sensitivity. In FETs, electrostatic gating of semiconducting channels upon target recognition leads to measurable changes in source-drain transconductance.^5,6^

For the detection of small molecules that have little to no charge, structure-switching aptamers can amplify the target-binding signal by rearranging the negatively charged oligonucleotide backbone and associated solution ions near the semiconductor surface.^5,7^ For this sensing mechanism, the net redistribution of the aptamer backbone upon target recognition drives the sensor response. To this point, characterization of the conformational dynamics of aptamers at FET surfaces upon interactions with small-molecule targets, is critical for analyzing the sensor response. Among the various biological targets explored with aptamer-FETs^5,8,9^, neurotransmitters such as dopamine and serotonin have generated high interest due to the importance of monitoring such small molecules to resolve brain function.^10–13^. To sense neurotransmitters *in vivo*, it is critical to consider the ionic content of cerebrospinal fluid (CSF), as solution ions influence oligonucleotide folding.^14–16^ In contrast to conventional buffers used for sensor testing (*e*.*g*., phosphate buffered saline), CSF contains millimolar (mM) concentrations of the divalent cations, Mg^2+^ and Ca^2+^. In particular, positively charged divalent cations can intercalate into the negatively charged DNA through electrostatic interactions, which can significantly influence the folding, stability, and overall conformation of G-quadruplex structures.^17^ Recent reports showed the influence of these divalent cations for dopamine aptamer-FETs in artificial CSF (aCSF).^18^ The presence of Mg^2+^ and Ca^2+^ on the dopamine aptamer-FETs manifested in an order-of-magnitude sensor signal enhancement. Divalent cations were hypothesized to induce a larger structure switching for the dopamine aptamer upon target capture, leading to a greater change in recorded signal. Alternatively, this effect was not observed for the serotonin aptamer. Herein, MD simulations were used to investigate and to corroborate the experimental findings between the two aptamers to gain insights into the sensing mechanism of aptamer-based FET sensors. In the MD simulations, the aptamers were constrained at the 5’ end to mimic covalently-tethered states with reduced degrees of molecular freedom on the sensor surface (details in SI-note#1). Structural analyses of the structure switching of the dopamine and serotonin aptamers were conducted in different electrolytes, namely aCSF and aCSF lacking Mg^2+^ and Ca^2+^, to mimic experimental conditions.^18^ Additionally, configurations where only one of the two divalent cations were removed from the aCSF were probed (see SI-note#2). During the MD simulations, the analytes were intentionally placed in various initial positions, to confirm target recognition to the same binding pocket.

According to experimental findings, the structure-switching behavior of the serotonin aptamer upon target binding was unchanged in different electrolytes^18^. From the MD simulations, we observed that in aCSF, the serotonin aptamer-target formed a stable complex due to the formation of four stable hydrogen bonds and electrostatic interactions (Pi-Anion, favourable acceptor-acceptor, and van der Waals, SI – note#3). The 3-D conformation obtained from the MD simulations pre- and post-serotonin recognition, depicted an elongation of approximately 3 nm of the serotonin aptamer backbone upon interaction with serotonin as shown in Fig. 1a. This elongation was previously observed experimentally using Förster resonance energy transfer (FRET) ^18^. In this study, we replicated the placement of donor and acceptor dyes used in FRET within the simulation to compare the 3-D structural rearrangement of the serotonin aptamer upon target capture. As schematically depicted in Fig. 1a, the measurements revealed that in the absence of serotonin, the separation between the two dyes was ∼1 nm, a distance conducive to FRET. However, in the presence of serotonin, this distance extended to ∼6 nm, leading to fluorescence quenching. Hence, the simulated aptamer conformation aligns with experimental FRET results. These findings reinforce the validity of the methodology used herein to extract aptamer 3-D conformations. We note that the conformations shown in this manuscript, represent the most frequent conformation extracted from the simulation trajectory. Other conformations are shown in SI-note#4). The serotonin aptamer MD simulations in aCSF *vs*. aCSF lacking divalent cations, revealed that the conformations of the unbound aptamer and the aptamer-serotonin complex are retained in both solutions, in agreement with prior FET measurements^18^ (SI – note#5).

**Fig. 1.**
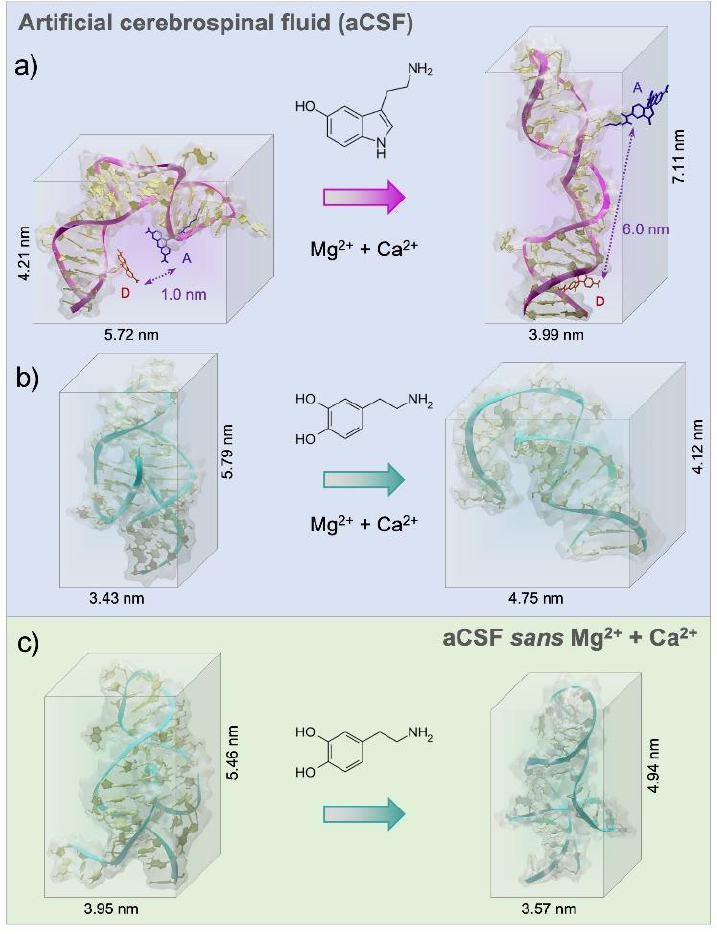
a) Extracted 3-D conformations from the molecular dynamic (MD) simulations run for 200 ns of a) the serotonin aptamer in the absence and presence of serotonin in artificial cerebrospinal fluid (aCSF) (details on the FRET distance simulation), (b) the dopamine aptamer before and post dopamine exposure in aCSF, and (c) the free dopamine aptamer and dopamine-bound complex in aCSF (*sans* Mg^2+^, Ca^2+^). For the dopamine aptamer, large conformational rearrangements occurred in the presence of divalent cations (Mg^2+^, Ca^2+^) and removal of these cations led to smaller structural rearrangements upon target capture. To improve the visibility, all water molecules and ions inside the simulation box were removed.

The dopamine aptamer MD simulations in aCSF revealed the presence of four stable hydrogen bonds, a salt bridge, Pi-Pi, and T-shape interactions for molecular recognition of dopamine (see SI– note#6). The analysis uncovered the binding pocket located within the asymmetric interior loop of the dopamine aptamer. Extracting the 3-D structures of the dopamine aptamer at the end of the MD simulations, we observed a compression of approximately 1.7 nm upon dopamine binding (Fig. 1b). However, in the absence of divalent cations, both the free aptamer and the dopamine-bound state exhibited altered conformations with respect to the previous cases (Fig. 1c). Differences in both the location of the binding pocket and the magnitude of conformational rearrangement were observed.

Individual effects of each divalent cation (Mg^2+^ and Ca^2+^) were then investigated to understand the cation size effect on the dopamine aptamer conformation. Visible differences in the target-induced conformational changes were observed for the dopamine aptamer in the absence of Ca^2+^ (Fig. 2a) *vs*. Mg^2+^ (Fig. 2b). Specifically, the dopamine aptamer without the larger cation, Ca^2+^, exhibited maximal structure switching, suggesting the importance of Mg^2+^ for folding. Smaller Mg^2+^ ions are effective at intercalating into G-quadruplex structures leading to stronger ion-DNA interactions and a more stable conformation.^19^

**Fig. 2.**
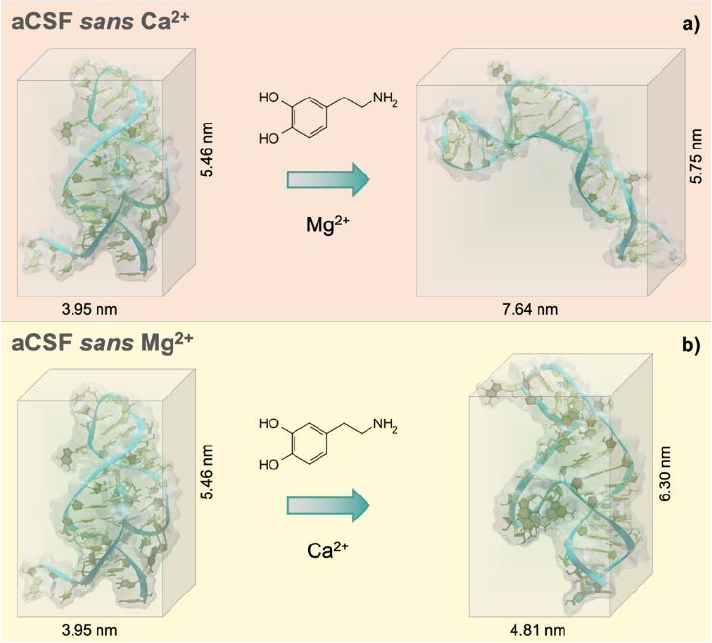
Extracted 3-D conformations from the molecular dynamic simulations of the dopamine aptamer and dopamine target in a) artificial cerebrospinal fluid (aCSF) lacking Ca^2+^, b) aCSF lacking Mg^2+^. While the starting conformation of the free dopamine aptamer remained unchanged regardless of the absence of one of the two divalent cations, the conformational change post-dopamine recognition was significantly altered based on the ionic species present. To improve the visibility, all water molecules and ions inside the simulation box were removed.

To gain an improved understanding on the influence of divalent cations on the behavior of free dopamine and serotonin aptamers, the formation of potential G-quadruplex structures was investigated. We observed exclusive formation of a G-quadruplex secondary structure within the dopamine aptamer in artificial cerebrospinal fluid (aCSF), as illustrated in Fig. S8. However, in the absence of either one or both of the divalent ions (Mg^2+^ and Ca^2+^) from the electrolyte, no G-quadruplexes were formed within the dopamine aptamer. In contrast, the serotonin aptamer, regardless of the electrolyte composition (aCSF, *sans* Mg^2+^, *sans* Ca^2+^, or lacking both Mg^2+^ and Ca^2+^), did not exhibit any G-quadruplex structure. This divergence in sensitivity to divalent cations between the dopamine and serotonin aptamers even for the starting conformations demonstrates the potential influence of ionic content on the secondary structure of aptamers.

The absence of both divalent cations resulted in a different dopamine-bound structure compared to the aptamer lacking solely Mg^2+^. This effect may be due to the combined presence of both cations leading to greater ion-ion interactions, which can modulate the distribution and ion availability near the aptamer and lead to distinct structural outcomes. Ion-ion interactions may influence the stability and folding propensity of G-quadruplexes, resulting in conformational states that differ from when each ion is present individually. Further, screening effects due to the cations may contribute to the differential conformational changes. Electrostatic interactions can shield or reduce the effective charges in specific DNA regions.^20^ Consequently, electrostatic repulsions between negatively charged areas may lead to different conformational landscapes for the aptamer. These screening effects may vary between Mg^2+^ and Ca^2+^ ions due to their differences in charge density and hydration effects.^20^ Taken together, these hypotheses emphasize the need to explore the roles of divalent cations in the conformational changes of each individual aptamer. Investigating the variations in charge distribution, ion-DNA and ion-ion interactions, and screening effects will provide valuable insights into the underlying mechanisms driving the observed differences in aptamer conformational dynamics.

The results obtained from the MD simulations can be compared to the experimental results reported in the literature on aptamer-modified FET biosensors.^18^ The conformational change of the negatively charged aptamer backbone induced by target recognition at the semiconductor surface, leads to charge rearrangement within or near the Debye length (λ_D_, ∼0.7 nm in physiological systems). While the Debye profile *vs*. screening distance is logarithmic in nature, for simplicity, a value of 0.7 nm was considered. Starting from the MD 3-D models, the number of negatively charged oligonucleotides that fit within this Debye length were calculated (Fig. S9, Table S2 – SI – note#7).

In the case of the serotonin aptamer, minimal differences were observed upon the removal of the divalent ions. In the absence of serotonin the number of nucleotides within λ_D_ was 6±2 for all cases. Following the serotonin-specific interaction, this number was decreased to 3±2 (Fig. S9). For the dopamine aptamer, upon removal of the divalent cations, the number of nucleotides within the λ_D_ for the free aptamer increases from 2±1 to 7±2. After binding to dopamine, the number of negatively charged nucleotides within λ_D_ changed in a varying range for the different cationic conditions (Fig. S9). In particular, for the aptamer in aCSF, the number of nucleotides within λ_D_ *increased*. This effect can be correlated to the experimental observations on FET surfaces. An increased density of negative charges due to the dopamine aptamer folding upon target capture in aCSF, caused electron depletion in the *n*-type channel, and led to a reduced current response. Conversely, when either Mg^2+^, Ca^2+^ or both cations were absent, the dopamine aptamer rearrangement led to electron accumulation in the FET semiconducting channel and a subsequent increase in measured current. The greater the redistribution of the negatively charged backbone (delta) within the λ_D_ post analyte binding, the higher the expected sensor response.^18^ These results offer significant insights into the operational mechanism of aptamer-modified biosensors. By delving into aptamer conformational dynamics at a molecular level, this research illuminates the underlying processes that drive aptamer-modified FET behavior.

Another established method to track DNA conformational changes and dynamics is circular dichroism (CD) spectroscopy.^18^ While CD is commonly conducted for polypeptides and proteins,^22^ aptamer-target interactions in solution can also be investigated. While CD shows spectral shifts representative of new motif formation due to rearranged base stacking and dipole moments in the molecule, it is impossible to extract the DNA molecular configuration, including target binding pocket and 3-D charge density distribution from the spectra. In this context, MD simulations can be used to extract the atomic coordinates of the DNA under investigation and these data can be used to calculate theoretical CD spectra. The CD spectra for the aptamers were calculated using DichroCalc^21^. Similar features were observed between experimental and calculated spectra, notably negative CD bands in the region 230-260 nm and positive CD bands in the region 250-290 nm (Fig. 3 a-c). Analysis of the computed CD spectra for the three cases of the dopamine aptamer (SI – note#8, Fig. S12 and Table S3) indicates that for each structure, different nucleic acid bases contribute to the CD bands. Bases are considered to contribute significantly if their configuration interaction coefficient (eigenvector matrix element) is greater than |0.5|. Results are reported for the three cases for the dopamine aptamer (Fig. 3d).

**Fig. 3.**
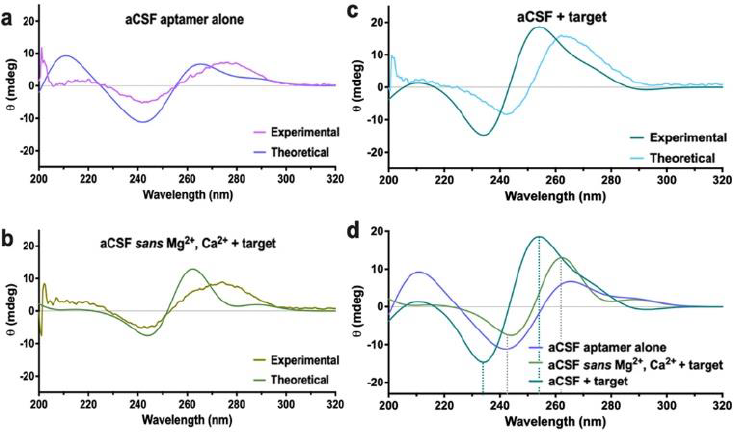
Circular dichroism (CD) spectra – comparison between experimental and calculated data for the dopamine aptamer a) free in artificial cerebrospinal fluid (aCSF) solution, b) in aCSF lacking Mg^2+^ and Ca^2+^ in the presence of dopamine, and c) in aCSF with dopamine. d) The calculated CD spectra for each condition are overlaid to illustrate the target-specific spectral shift observed in the experimental CD data.

Computational tools indicated that for the target-bound dopamine aptamer in aCSF, the negative band at 234 nm is due to bases 8A, 14A, 15A and 35A, and the positive band at 254 nm due to bases 19T, 21G, 22T 25G and 33G. The unbound dopamine aptamer in aCSF has bases 3A, 9A, 13A and 15A contributing to the negative band at 242 nm, with bases 11G, 20G, 32G, 35G, 37G, 40G and 41G contributing to the positive band at 265 nm. For the dopamine-bound aptamer in aCSF *sans* cations, the negative band at 244 nm is due to bases 8A, 10T, 11T, 14A, 15A, 27A and 39A, and the positive band at 262 nm due to bases 5G, 25G, 31G and 36G. For each nucleobase, the local ππ* transitons responsible for the rotational strengths are at wavelengths 241.7 nm for A, 248.0 nm for T, 254.1 nm for T and 261.1 nm for G. These analyses elucidate the different nucleotides (and aptamer binding sites) influenced by the different electrolytes and can be the basis for further investigations combining CD and MD simulations in aptamer molecules.

In summary, this work describes the use of MD simulations to investigate the behaviour of aptamers in the presence of neurotransmitters in different ionic environments. Specifically, the dopamine and serotonin aptamers were studied, with a focus on understanding the effect of divalent cations present in brain fluid. We demonstrated how divalent cations influenced the structure-switching behaviour of the dopamine aptamer while minimally perturbing the serotonin aptamer. The 3-D structures obtained from MD simulations were used to elucidate the working mechanism of FET-biosensors. Moreover, the role of divalent cations in stabilizing the aptamer conformations and preserving the binding pockets is highlighted, shedding light on the importance of the microenvironment in influencing aptamer-target interactions.

This study not only theoretically analyzed the dynamic conformations of aptamers but also correlated these findings with experimental measurements of aptamer-modified FETs, particularly in terms of signal magnitude and directionality. The findings herein, offer a comprehensive molecular perspective of the structure-switching dynamics at sensor surfaces, which supports existing hypotheses on the operational dynamics of FET-based biosensors.

