## Supporting Information for "Theoretical Analysis of Divalent Cation Effects on Aptamer Recognition of Neurotransmitter Targets"

### Supporting note #1 – Aptamers sequences and technical details on MD Simulations

The molecular dynamics (MD) simulations investigating the interaction between dopamine/serotonin and the aptamer were conducted using Gromacs software, employing the AMBER99SB-BSC1 force field. Initially, the aptamer was positioned at the center of a water box, which was appropriately sized based on the dimensions of the different aptamers. The dopamine or serotonin targets/analytes were then introduced into the box at various initial positions, maintaining a distance of 3 nm between the box and the aptamer surface. Subsequently, water molecules following the TIP3P model and ions to maintain charge neutrality were added to the system. The topology files for the analytes were generated using AmberTools within Gromacs. To achieve energy minimization, the steepest descent algorithm was applied, considering periodic boundary conditions (PBC) in all directions. The equilibration of the systems was carried out under constant number (N), volume (V), and temperature (T) conditions (NVT ensemble) and constant number (N), pressure (P), and temperature (T) conditions (NPT ensemble), with a time step of 2 fs, lasting 10 ns. Subsequently, a 300 ns simulation was performed under a pressure of 1.01 atm and a temperature of 294 K. Additionally, the equilibrated system was simulated in the NPT ensemble for an additional 10 ns.

The selection of five conformations was a critical step in our study, following a defined pipeline (Scheme 1). We initiated by determining the DNA aptamer's secondary structure using mFold software (http://www.unafold.org/mfold/applications/dna-folding-form.php). Then, the 3D models based on the aptamer sequence and secondary structure were generated using the 3dRNA/DNA server. Afterwards, the structures underwent energy minimization, equilibrations (NVP and NPV), and a 50 ns simulation at 2 picosecond intervals. Frames from the equilibration phase were discarded based on RMSD plots to retain only relevant data. Trajectories were analyzed using the Gromacs g-cluster tool, clustering structures with a 2 Å RMSD cutoff, yielding 20 different clusters. To ensure thorough exploration, we conducted 20 independent simulations to cover various energy and conformational spaces. The resulting trajectories were concatenated into a single pseudo-trajectory, and further clustering identified the five largest clusters for subsequent docking analysis. We also employed free-energy landscape (FEL) and principal component analysis (PCA) to assess the study's robustness and conformational exploration. Ultimately, cluster centers, representing metastable states, served as the basis for our molecular docking analysis with the corresponding ligands. This comprehensive approach guaranteed diversity in conformations and strengthened the validity of our findings regarding aptamer-ligand interactions.

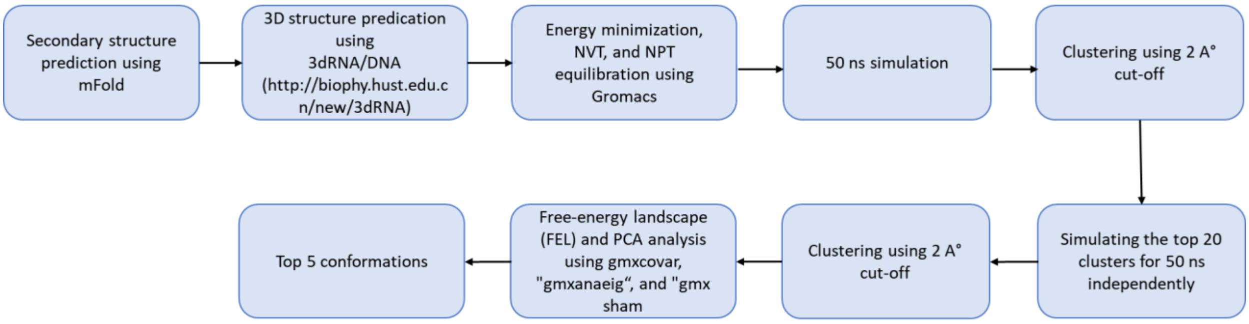

Scheme 1. Workflow of the pipeline used to analyze aptamer–small-molecule interactions.

Single-stranded DNA sequences were used in this work:

_Dopamine aptamer sequence:_

CGACGCCAGTTTGAAGGTTGGTTCGCAGGTGTGGAGTGACGTCG

_Serotonin aptamer sequence:_

CGACTGGTAGGCAGATAGGGGAAGCTGATTCGATGCGTGGGTCG

**Supporting note #2 – Electrolytes compositions used in the MD Simulations**

| aCSF | | aCSF Sans $Mg^{+2}$ | | aCSF Sans $Ca^{2+}$ | | aCSF Sans $Mg^{+2}/Ca^{2+}$ | |
|---|---|---|---|---|---|---|---|
| Salt | Conc. (mM) | Salt | Conc. (mM) | Salt | Conc. (mM) | Salt | Conc. (mM) |
| NaCl | 147 | NaCl | 147 | NaCl | 147 | NaCl | 147 |
| KCl | 3.5 | KCl | 3.5 | KCl | 3.5 | KCl | 3.5 |
| $MgCl_2$ | 1.2 | $CaCl_2$ | 1.0 | $MgCl_2$ | 1.2 | | |
| $CaCl_2$ | 1.0 | | | | | | |

**Table S1**: Composition of the electrolytes used during the MD simulations.

**Supporting note #3 – Serotonin Aptamer – Serotonin interaction**

MD simulations provide a detailed analysis of the binding phenomena at each time step, revealing the interactions between individual nucleobases of the aptamer and the neurotransmitters, as well as the resulting noncovalent chemical interactions for molecular recognition.

Notably, hydrogen bonds were formed between the hydrogen atoms of serotonin and the aptamer's nucleotides G19, G20, T26, and G27, at precise distances of 2.12 Å, 2.59 Å, 1.89 Å, and 2.79 Å, respectively. Additionally, electrostatic interactions were discerned between the nucleotide A17 and the amine group of serotonin, and between the nucleotide G18 and the primary amine of serotonin, further underscoring the exquisite nature of the intermolecular interactions (Fig. S5).

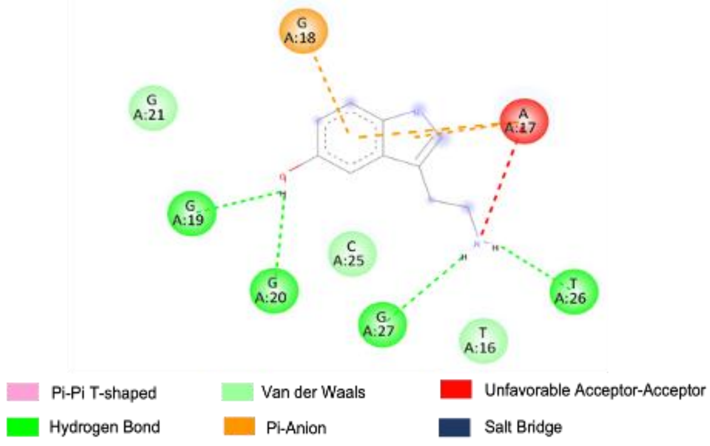

**Fig. S1.** Interactions observed in the two-dimensional representations obtained from the molecular dynamics simulations are shown for the serotonin aptamer-serotonin interactions in aCSF. The aptamer nucleotides are depicted as colored spheres surrounding the molecular structures. Various types of interactions are indicated by different lines: green dashed lines represent hydrogen bonds, pink lines indicate Pi-Pi T-shaped interactions, orange lines represent Pi-anion interactions, red lines indicate unfavorable acceptor-acceptor interactions, and blue lines represent salt bridges. Nucleotides highlighted in light green without dashed line connections are involved in Van der Waals interactions with the analyte.

### Supporting note #4 – Energy landscape

To gain insights into the crucial dynamics and structural variations within the aptamers, principal component analysis (PCA) was conducted on the backbone atoms of each aptamer. The analysis was based on the last 50 ns trajectories of the simulations. The "gmxcovar" tool in Gromacs was utilized to generate eigenvectors then a 2D projection and Energy Landscape (FEL) plots were constructed using the "gmxanaeig" and "gmx sham" utilities in Gromacs, respectively. In these FEL diagrams, we projected the first and second principal components (PC1 and PC2) to generate 2D representations for the aptamers systems, shown in Figures S1a–c. Notably, these FEL plots revealed a predominant low-energy region (local minimum) for each of the dopamine and serotonin states. This observation strongly suggests that the backbone atoms within each system tend to adopt a dominant low-energy structure. This behavior is likely a result of the clustering process that was previously conducted, which enhanced the probability of extracting the dominant structure. In summary, our findings indicate that any structural variations in the aptamers are likely confined to specific local structural elements rather than being widespread throughout the overall aptamer structure.

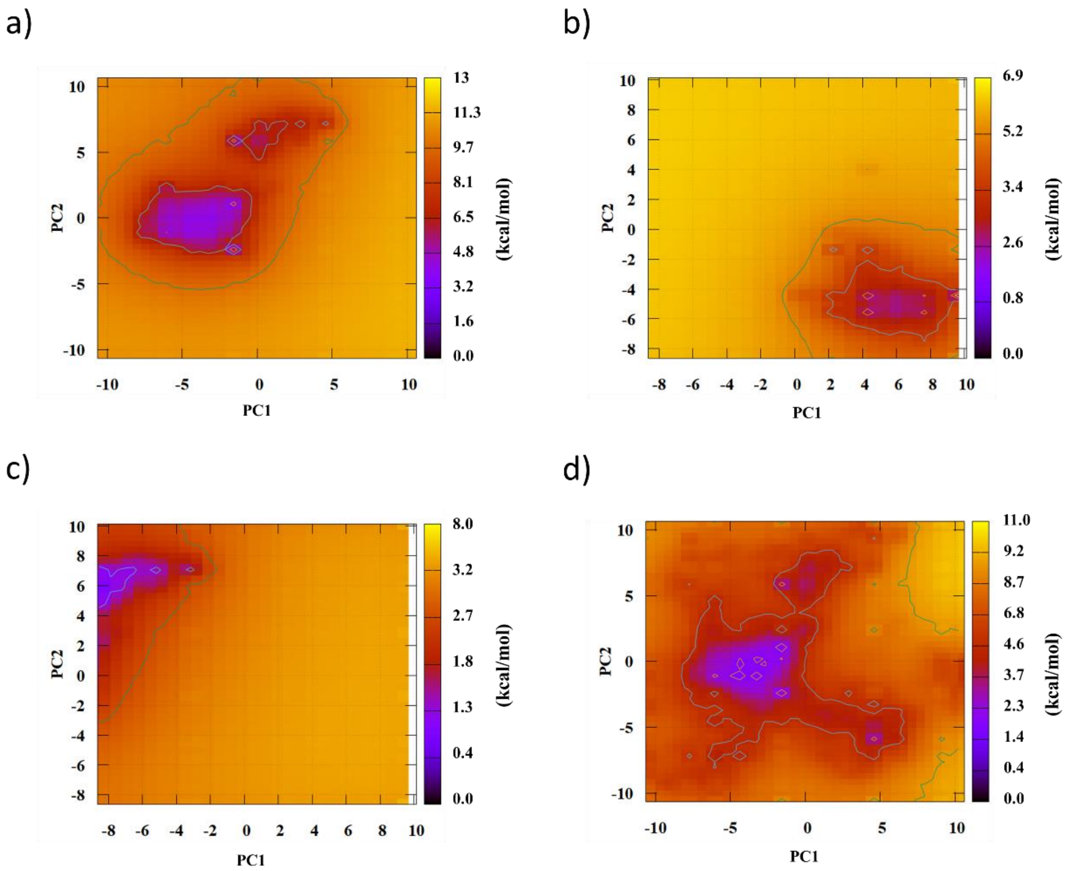

**Fig. S2.** Free energy landscape (FEL) contour plots of a) serotonin aptamer-serotonin complex in artificial cerebrospinal fluid (aCSF), b) dopamine aptamer-dopamine complex in aCSF, c) dopamine aptamer-dopamine complex in aCSF (Sans $Ca^{2+}$), d) dopamine aptamer-dopamine complex in aCSF (Sans Mg 2+).

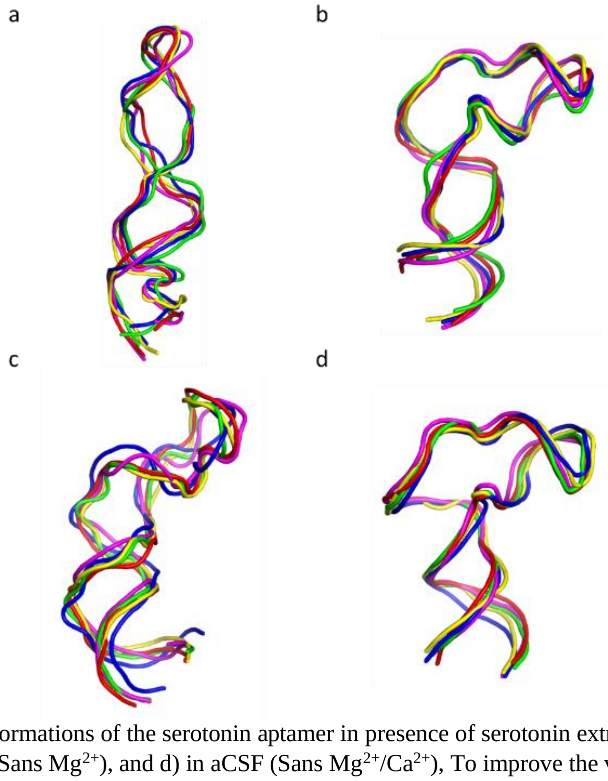

**Fig. S3.** a) Top five frequent 3-D conformations of the serotonin aptamer in presence of serotonin extracted after clustering a) in aCSF, b) aCSF (Sans $Ca^{2+}$), c) aCSF (Sans $Mg^{2+}$), and d) in aCSF (Sans $Mg^{2+}$/$Ca^{2+}$), To improve the visibility, all water molecules and ions inside the simulation box were removed.

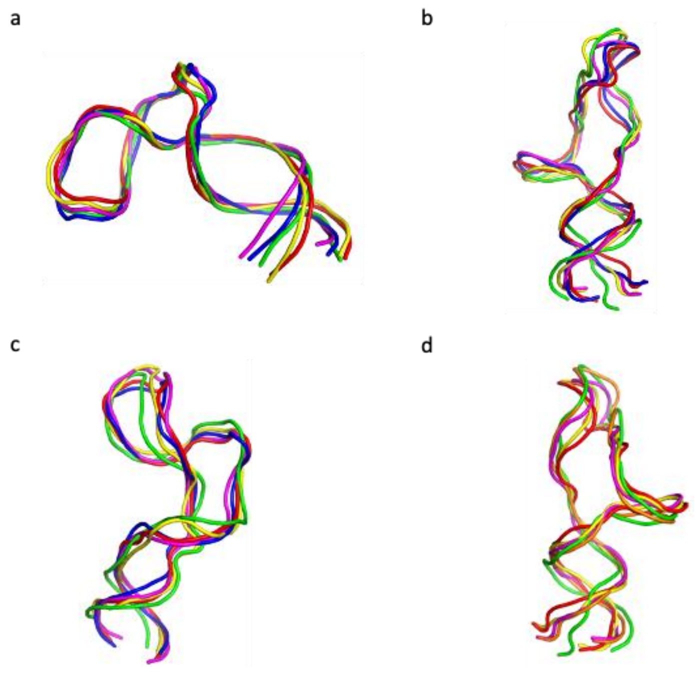

**Fig. S4.** a) Top most frequent 3-D conformations of the dopamine aptamer in presence of dopamine extracted after clustering a) in aCSF, b) aCSF (Sans $Ca^{2+}$), c) aCSF (Sans $Mg^{2+}$), and d) in aCSF (Sans $Mg^{2+}/Ca^{2+}$), To improve the visibility, all water molecules and ions inside the simulation box were removed.

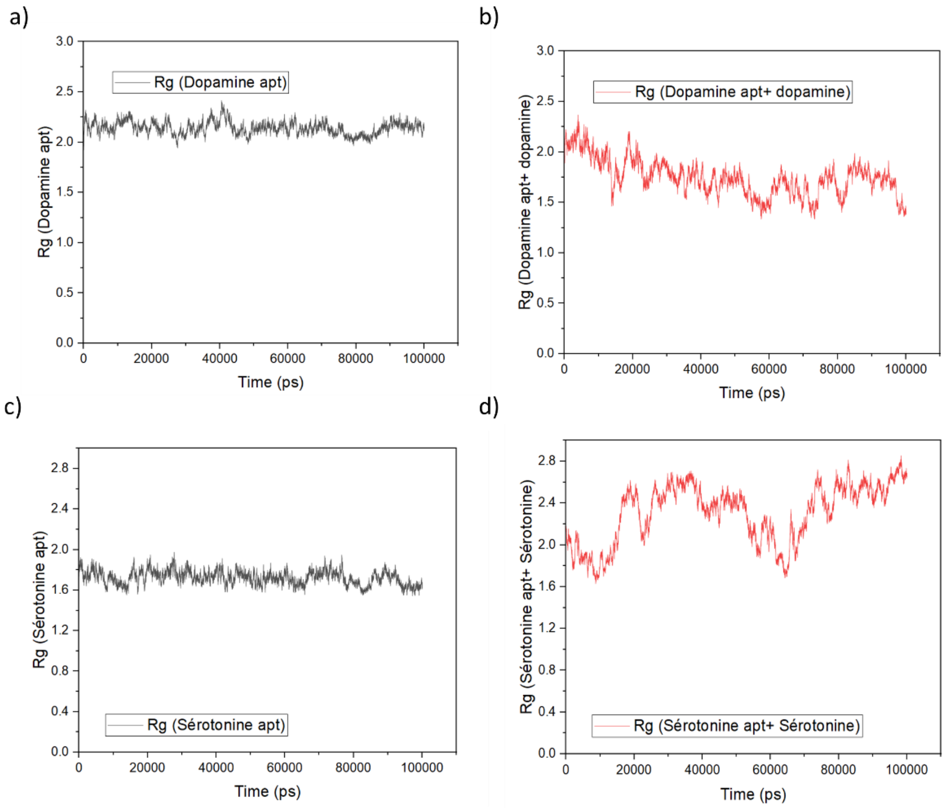

**Fig. S5.** Rg values of a) dopamine aptamer in aCSF, b) dopamine aptamer-dopamine in aCSF, c) serotonin aptamer in aCSF, d) serotonin aptamer-serotonin in aCSF.

### Supporting note #5 - Individual effects of each divalent cation ($Mg^{2+}$ and $Ca^{2+}$) on Serotonin Aptamer

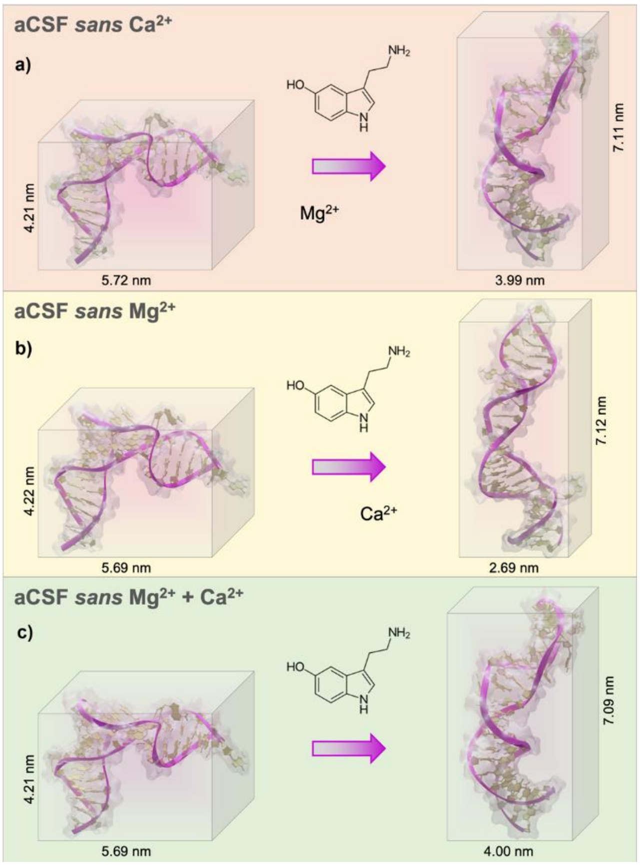

**Fig. S6.** a) Extracted 3-D conformations of the serotonin aptamer and serotonin molecule in aCSF (Sans $Ca^{2+}$), b) Extracted 3-D conformations of the serotonin aptamer and serotonin molecule in aCSF (Sans $Mg^{2+}$), c) Extracted 3-D conformations of the serotonin aptamer and serotonin molecule in in aCSF (Sans $Mg^{2+}$/$Ca^{2+}$), To improve the visibility, all water molecules and ions inside the simulation box were removed.

### Supporting note #6 – Dopamine Aptamer – Dopamine interaction

In aCSF, hydrogen bonds were formed between the hydrogen atoms of dopamine and the aptamer nucleobases A28, A30, and G33, at precise distances of 3.12 Å, 4.42 Å, and 3.08 Å, respectively. Moreover, the primary amine (protonated at physiological pH) of dopamine interacted via electrostatic interactions with the nucleobases G33 (Pi-cation), A35 (salt-bridge), and A27 (Pi-anion), exhibiting an exquisite network of intermolecular interactions. Notably, a Pi-Pi T-shaped interaction between G29 and dopamine's aromatic group was discernible, further corroborating the strength of the binding between the aptamer and the dopamine molecule (Fig. S7a).

In the absence of divalent cations, both in the case of the free aptamer and the aptamer interacting with dopamine (Fig. S7b). The interaction revealed the formation of four stable hydrogen bonds and electrostatic interactions (Pi-Anion, Pi-Pi-T-shaped, salt-bridge). Hydrogen bonds were observed between dopamine's hydrogens and aptamer nucleobases T12, T11, and A35, with distances of 3.12 Å, 4.42 Å, and 3.08 Å, respectively. Electrostatic interactions were evident between T32 and G33 (Pi-cation), the primary amine of dopamine and A35 (salt-bridge), and Pi-anion between A27 and the amine of dopamine. Additionally, a Pi-Pi T-shaped interaction between G34 and the amine of dopamine was also present.

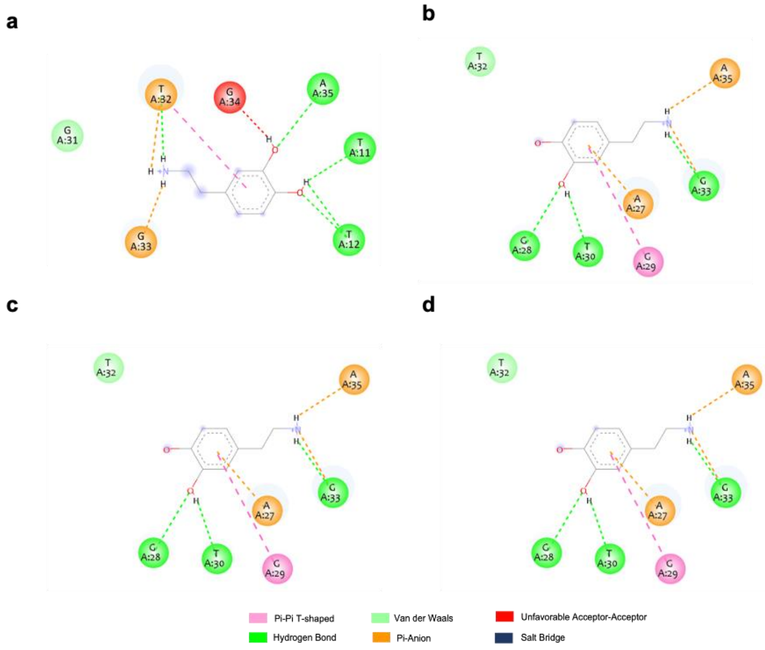

**Fig. S7.** Interactions observed in the two-dimensional representations obtained from the molecular dynamics simulation are shown for a) the dopamine aptamer-dopamine interactions in aCSF, b) in aCSF (Sans $Mg^{2+}$/$Ca^{2+}$), c) in aCSF (Sans $Mg^{2+}$), d) in aCSF (Sans $Ca^{2+}$). The aptamer nucleotides are depicted as colored spheres surrounding the molecular structures. Various types of interactions are indicated by different lines: green dashed lines represent hydrogen bonds, pink lines indicate Pi-Pi T-shaped interactions, orange lines represent Pi-anion interactions, red lines indicate unfavorable acceptor-acceptor interactions, and blue lines represent salt bridges. Nucleotides highlighted in light green without dashed line connections are involved in Van der Waals interactions with the analyte.

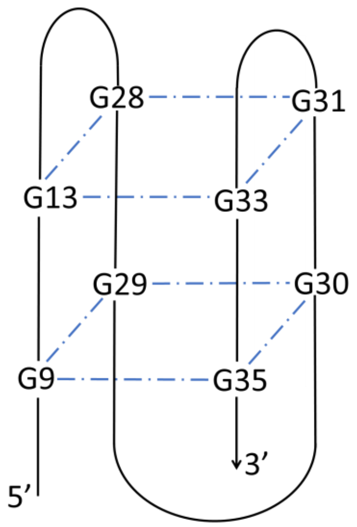

**Fig. S8**. Schematic showing h-bond interactions between G bases within the dopamine aptamer in aCSF.

### Supporting note #7 - Number of charges within the Debye length

*To calculate the number of nucleotides within the Debye length for each case, first the root mean square fluctuation (RMSF) values of each of the bases was calculated. Then, the fluctuation of the bases during the trajectory near the Debye length were taken into account, and the number of bases within the Debye length was calculated. Figures S10 and S11 show the RMSF values vs. residue for each of the cases. The residues stable within the Debye length are highlighted in blue and the residues that were fluctuating in and out of the Debye length are highlighted in red.*

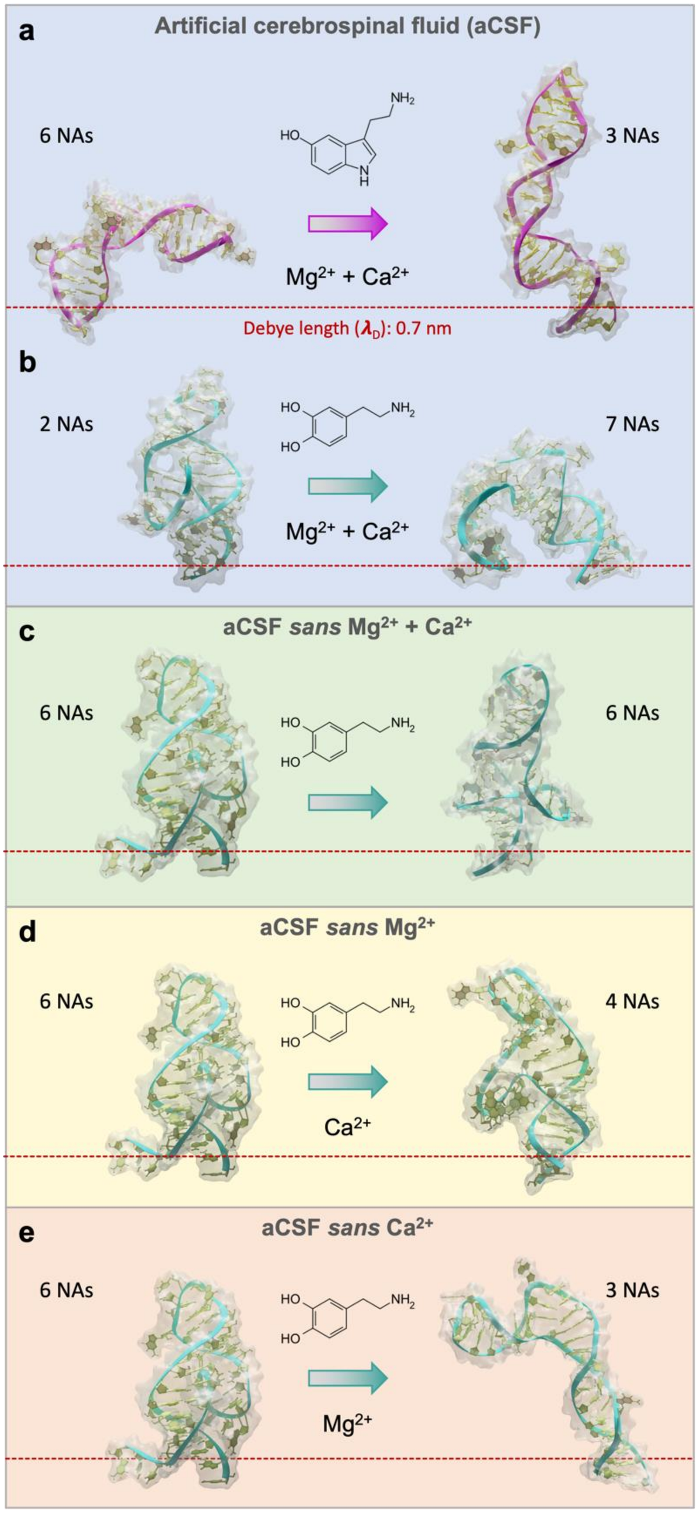

**Fig. S9.** Number of nucleic acids from the aptamers within the Debye length before (left side) and after interaction with the target molecule (right side). (a) case for Serotonin Aptamer; (b-e) cases for Dopamine aptamer.

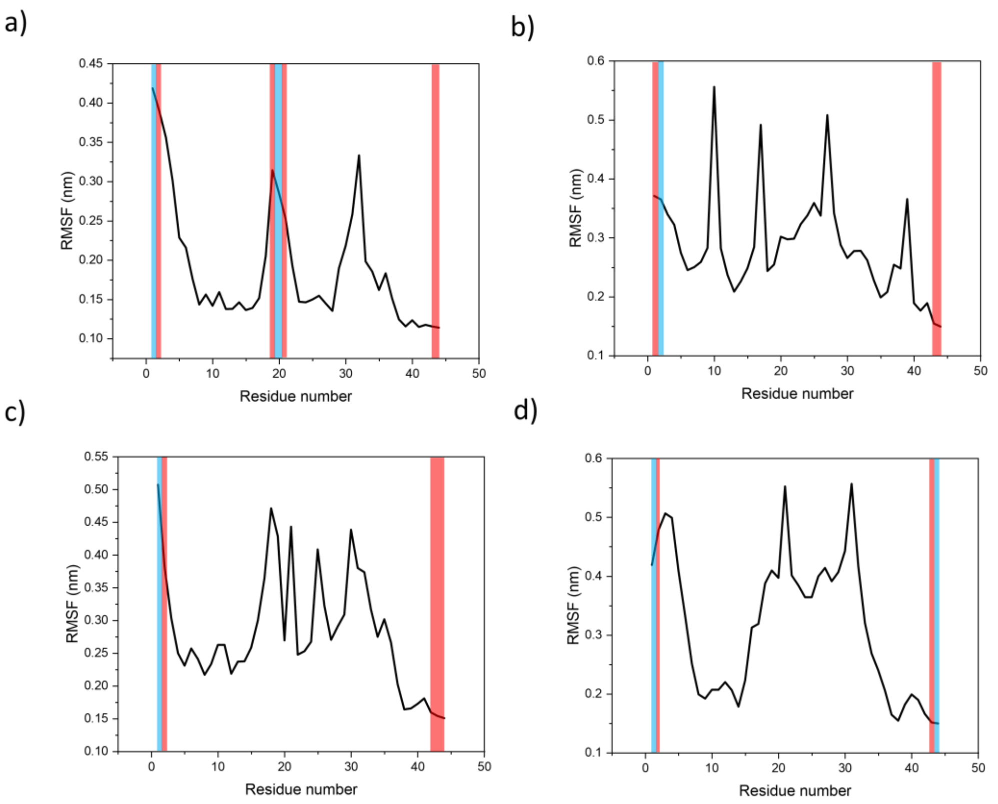

**Fig. S10**. Root mean square fluctuation (RMSF) graphs representing molecular dynamic simulations for 100 ns of a) dopamine aptamer-dopamine in artificial cerebrospinal fluid (aCSF), b) dopamine aptamer-dopamine in aCSF (Sans $Ca^{2+}$), c) dopamine aptamer-dopamine in aCSF (Sans $Mg^{2+}$), and d) dopamine aptamer-dopamine in aCSF (Sans $Mg^{2+}$/$Ca^{2+}$). The x-axis represents the residue number and y-axis represents the RMSF value (nm). Color scheme: blue (bases stable within the Debye length during the trajectory), red (bases fluctuating in and out of the Debye length).

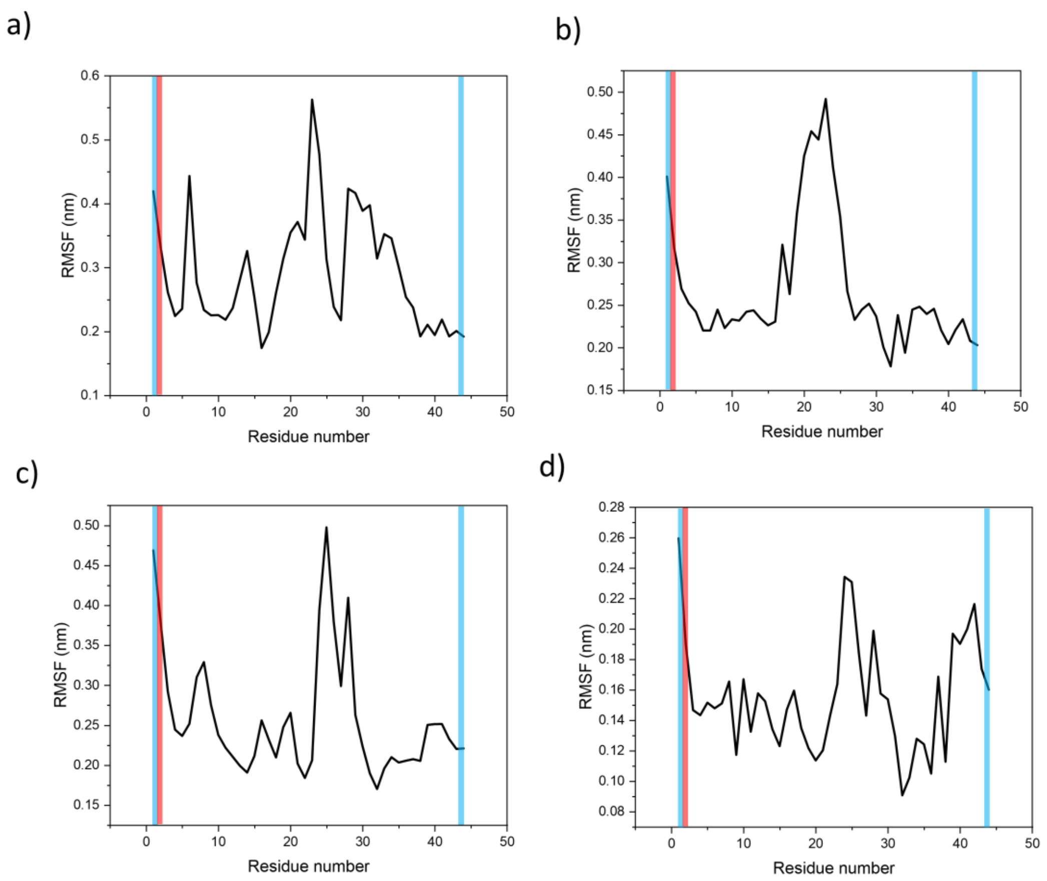

**Fig. S11.** Root mean square fluctuation (RMSF) graph representing molecular dynamic simulations for 100 ns of a) serotonin aptamer-serotonin in artificial cerebrospinal fluid (aCSF), b) serotonin aptamer-serotonin in aCSF (Sans $Ca^{2+}$), c) serotonin aptamer-serotonin in aCSF (Sans $Mg^{2+}$), and d) serotonin aptamer-serotonin in aCSF (Sans $Mg^{2+}$/$Ca^{2+}$). The x-axis represents the residue number and the y-axis represents the RMSF value (nm). Color scheme: blue highlight (bases stable within the Debye length during the trajectory), red highlight (bases fluctuating in and out of the Debye length).

| Aptamer | Before Target Interaction | After Target Interaction |
|---|---|---|
| Serotonin (aCSF) | 6 ± 2 | 3 ± 1 |
| Serotonin (aCSF sans $Mg^{2+}$ and $Ca^{2+}$) | 6 ± 2 | 3 ± 1 |
| Dopamine (aCSF) | 2 ± 1 | 7 ± 2 |
| Dopamine (aCSF sans $Mg^{2+}$ and $Ca^{2+}$) | 6 ± 2 | 6 ± 1 |
| Dopamine (aCSF sans $Mg^{2+}$) | 6 ± 2 | 4 ± 1 |
| Dopamine (aCSF sans $Ca^{2+}$) | 6 ± 2 | 3 ± 2 |

**Table S2:** Summary of the nucleotides within the Debye length obtained from Fig. 3.

**Supporting note #8 - DichroCalc computed CD for Dopamine Aptamer**

DichroCalc [1] was employed to calculate CD spectra for the dopamine aptamer. Unpublished parameter sets ADE-FUL3, GUAWFUL3, CYT-FUL3 and URA-FUL3 were used to describe two local transitions on each of the nucleic acid bases. Gaussian functions of full-width half-maximum (FWHM) 12.5 nm were fitted to the computed rotational strengths to yield CD spectra. There are a total of 44 residues in each of the PDB files containing the atomic coordinates for the aptamers.

Table S3 displays an analysis of the local group transitions that contribute to the computed rotational strength that gives rise to CD bands at a given wavelength for the dopamine aptamers. Contributing transitions to the rotational strength are tabulated if they have an eigenvector matrix element greater than |0.2|. The contributing transitions also include those transitions that couple (interact) with the dominant group transition considered responsible for the rotational strength. Fig. S12 shows the computed rotational strengths and the CD spectra in the wavelength range 190 to 320 nm. For the computed CD bands of the DNA aptamers in the near UV region of the spectrum, the first local electronic transition on the nucleic acid bases is predicted to be responsible for the rotary strength.

| CD band (nm) | Rotational strength energy ($cm^{-1}$) | Rotational strength wavelength (nm) | Rotational strength (DBM) | Residue, transition (nm), eigenvector matrix element |
|---|---|---|---|---|
| **aCSF + target** | | | | |
| 234 | 41780 | 239.35 | -1.169 | 8A, 241.68, 0.537;<br>35A, 241.68, 0.513;<br>10T, 247.97, 0.378;<br>11T, 247.97, 0.294;<br>37T, 247.97, 0.351 |
| | 41915 | 238.58 | -0.206 | 14A, 241.68, 0.562;<br>15A, 241.68, 0.719;<br>19T, 247.97, 0.207;<br>22T, 247.97, 0.241 |
| 254 | 38161 | 262.05 | 0.547 | 17G, 262.12, 0.423;<br>21G, 262.12, 0.639;<br>25G, 262.12, -0.211;<br>33G, 262.12, -0.520 |
| | 38214 | 261.68 | 0.226 | 13G, 262.12, 0.284;<br>25G, 262.12, 0.830;<br>28G, 262.12, 0.232;<br>33G, 262.12, -0.332 |
| | 39770 | 251.44 | 0.191 | 19T, 247.97, -0.666;<br>22T, 247.97, 0.729 |
| **aCSF no target** | | | | |
| 242 | 41383 | 241.64 | -0.761 | 9A, 241.68, 0.575;<br>13A, 241.68, 0.778 |
| | 41389 | 241.61 | 1.197 | 9A, 241.68, -0.809;<br>13A, 241.68, 0.524 |
| | 41426 | 241.40 | -0.933 | 14A, 241.68, 0.310;<br>15A, 241.68, 0.850;<br>33A, 241.68, 0.377 |
| | 41503 | 240.94 | -0.126 | 3A, 241.68, 0.961;<br>2G, 261.12, 0.232 |
| 265 | 37606 | 265.91 | 0.155 | 40G, 261.12, -0.576;<br>41G, 261.12, 0.773 |
| | 37664 | 265.51 | -0.100 | 11G, 261.12, -0.644;<br>35G, 261.12, 0.513;<br>37G, 261.12, -0.511 |
| | 37764 | 264.80 | 0.211 | 20G, 261.12, -0.726; |

| | | | | |
|---|---|---|---|---|
| | | | | 21G, 261.12, 0.416; 32G, 261.12, -0.541 |
| **aCSF sans cations + dopamine** | | | | |
| 244 | 39979 | 250.13 | -0.313 | 10T, 254.07, 0.674; 11T, 254.07, 0.688 |
| | 41274 | 242.29 | 0.149 | 14A, 241.68, -0.668; 15A, 241.68, 0.730 |
| | 41358 | 241.79 | -0.146 | 8A, 241.68, 0.901; 39A, 241.68, -0.420 |
| | 41378 | 241.67 | 0.181 | 27A, 241.68, 0.980 |
| | 41479 | 241.09 | 0.154 | 14A, 241.68, 0.416; 39A, 241.68, 0.891 |
| | 41569 | 240.56 | -0.367 | 27A, 241.68, 0.718; 39A, 241.68, 0.667 |
| 262 | 38129 | 262.27 | 0.189 | 25G, 261.12, -0.494; 31G, 261.12, -0.212; 36G, 261.12, 0.793 |
| | 38139 | 262.20 | -0.847 | 25G, 261.12, 0.499; 31G, 261.12, 0.606; 34G, 261.12, 0.235; 36G, 261.12, 0.478 |
| | 39200 | 261.78 | 1.144 | 25G, 261.12, 0.583; 31G, 261.12, -0.608; 33G, 261.12, 0.286; 34G, 261.12, 0.297 |
| | 38241 | 261.50 | -0.385 | 42T, 254.07, -0.219; 5G, 261.12, 0.914 |

**Table S3.** Analysis of the local group transitions that contribute to rotational strength and the CD bands at a given wavelength for the dopamine aptamers. Rotational strengths considered if greater or equal to |0.1| DBM. Local group transitions with eigenvector matrix elements greater than |0.2|.

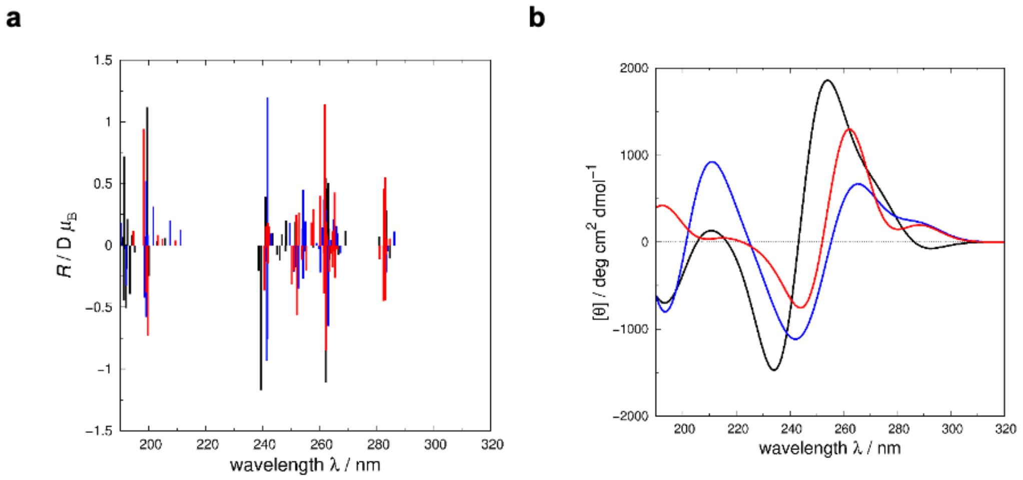

**Fig. S12.** a) Rotational strength and b) CD spectra, for the three dopamine aptamers aCSF + target (black line), aCSF no target (blue line) and aCSF sans cation + dopamine (red line). Gaussian functions of full-width half-maximum (FWHM) 12.5 nm were fitted to the computed rotational strengths to yield the CD spectra.
